# Unraveling cryptic diversity: Genomic approaches to study the taxonomy and evolution of Woolly-necked storks using museum specimens

**DOI:** 10.1101/2023.11.22.568311

**Authors:** Prashant Ghimire, Catalina Palacios, Jeremiah Tremble, Sangeet Lamichhaney

## Abstract

The availability of large-scale genomics data for current research in evolutionary biology has enabled a comprehensive examination of the intricate interplay between diverse evolutionary processes associated with speciation. Despite these advancements, the genomic basis of taxonomic classification remains challenging in many species, one such example being the Woolly-necked storks *(Ciconia sps.)*. The Woolly-necked storks are distributed in Asia and Africa with a taxonomic classification *(C. episcopus and C. microcelis)* that has been a matter of contention and ambiguity. Asian and African Woollynecks were just recently recognized as different species based on their morphological differences, however genetic/genomic studies on Woolly-necked storks are lacking. In this study, we have used ∼70-year-old museum samples to explore the taxonomy and evolution of the Woolly-necked storks. We used a whole-genome sequencing strategy and generated 13.5 million single nucleotide polymorphisms (SNPs) that were polymorphic among populations of Asian and African Woollynecks. Our study has revealed that Asian and African Woollyneck are genetically distinct, consistent with the current taxonomic classification based on morphological features. However, we also found a high genetic divergence between the Asian subspecies *C. e. neglecta* and *C. e. episcopus* suggesting this classification harbors cryptic diversity that requires a detailed examination to explore processes of ongoing speciation. Because taxonomic classification directly impacts conservation efforts, and there is evidence of declining populations of Asian Woollynecks in Southeast Asia, our results suggest populations-scale studies are urgent to determine the genetic, ecological, and phylogenetic diversity of these birds. Moreover, our study also provides historical genomic resources to examine genomic signatures of local adaptation associated with the distribution, ecology, and evolution of African and Asian Woollynecks.

## Introduction

Population divergence and speciation are still central subjects in evolutionary biology, although Darwin sparked the discussion one and a half centuries ago (Darwin 1859). Population divergence refers to the process by which populations accumulate genetic and phenotypic differences over time through factors including natural selection, genetic drift, mutation, and gene flow (Schluter 2001). Speciation refers to the formation of new species which occurs when populations become stable independent lineages, with specific phenotypic and genetic traits, and are reproductively isolated from one another (Mayr 1982; De Queiroz 2007). The relationship between population divergence and speciation is not always clear. Although population divergence may lead to the formation of new species, speciation may occur rapidly between low-divergent populations due to natural selection. Also, some species may maintain high levels of population divergence without resulting in speciation as in the case of generalist species with a wide distribution range (Engelbrecht et al. 2020; Liu et al. 2020).

Modern research in evolutionary biology has provided new insights into the mechanisms underlying population divergence and speciation (Bolnick et al. 2023). Advances in genomics and molecular biology have allowed scientists to examine the genetic basis of speciation and identify genes involved in reproductive isolation and divergence (Coyne 1992; Schluter 2009; Sobel et al. 2010). Additionally, studies using model organisms and comparative genomics have shed light on the different modes and rates of speciation (Gagnaire 2020; Genereux et al. 2020). With the availability of large-scale genomics data, characterization of the dynamic interplay of diverse evolutionary processes such as gene flow, mutation, recombination, drift, and selection in shaping the genomic landscape of speciation has been increasingly feasible (Ravinet et al. 2017; Campbell et al. 2018). Moreover, the use of museum specimens for genomics research has gained popularity in recent years as advancements in high-throughput genome sequencing technologies have made it possible to obtain valuable genomic data even from low-quality or historical biological samples (Card et al. 2021). However, because of the rapid decline of diversity across taxa, we may be losing diversity and species even before being able to characterize them (Dirzo and Raven 2003). This is particularly problematic in groups with cryptic diversity where distinct morphological traits are limited, and taxonomic status may not reflect the true genetic diversity.

Traditionally, species have been defined based on distinct, nonoverlapping morphological characteristics observed in individuals from geographically separated populations (Mayr 1999). Later, the availability of large-scale genomic data has helped in the task of defining species because genomic data adds resolution (millions of loci across the genome), the states of the characters (alleles) are easier to define, and generating genomic data has become increasable easy (Felsenstein 1988; Goodwin et al. 2016). However, species delimitation, the accurate determination of the species boundaries, remains challenging (Smith and Carstens 2022),for example: (a) species may display continuous or overlapping morphological variation patterns rather than clear differences (Grant 2014; Lamichhaney et al. 2015), (b) species may exhibit morphological variation that is not yet recognized or described due to limited sampling or lack of taxonomic expertise (National Academies of Sciences, Engineering, and Medicine and National Academies of Sciences, Engineering, and Medicine 2019), (c) there may be cryptic species that may be distinct in physiological, behavioral, genetic or other less conspicuous traits difficult to differentiate using traditional approaches alone (Bickford et al. 2007), (d) individuals within a species can exhibit phenotypic plasticity, presenting different morphs based on environmental conditions (Forsman 2015) (e) interspecific hybridization, where individuals from different species mate and produce offspring, blur the boundaries between species (Abbott et al. 2013) or (f) lack of enough genetic studies to support species characterization (Rannala 2015). Species delimitation has been a controversial topic in systematics (Zapata and Jiménez 2012) and our understanding of species diversity is constantly evolving. Currently, it is recognized that different lines of evidence are required to define a new species (Padial et al. 2010).

The genetic and phenotypic differences among populations are commonly shaped by the environment (Bonaccorso et al. 2006; Hernández-Romero et al. 2018). Geography plays a major role in separating well-connected populations reducing gene flow in favor of divergence and reproductive isolation (Coyne 1992). Related taxa spanning continents undergo disparate influences, including dynamic landscape alterations, climatic variations, seasonal shifts, and ecological interactions. Consequently, they face a diverse array of selective pressures, contributing to the observed variations in their characteristics and adaptations (Keith et al. 2013; Craw et al. 2016; Pellissier et al. 2018). Studies have indicated that the divergence time of sister clades located on different continents aligns with continental break-ups, providing additional evidence supporting the role of plate tectonics and its impact on biodiversity (Carney and Dick 2000; McIntyre et al. 2017). The separation of South America from Africa and Australia, for example, drove the evolution of unique species such as marsupials and monotremes (Long 2017). The collision of tectonic plates has also played a pivotal role in the dispersion of organisms across continents and has contributed to their diversification. An illustrative example is the Indian plate, which, upon breaking away from the Gondwanan landmass, facilitated biological dispersal along the Indian Ocean, e.g. Natatanuran frogs (Yuan et al. 2019). Similarly, the creation of islands and their dynamic processes foster alterations in the diversity of organisms inhabiting them. This phenomenon is evident in current hotspots of endemic biodiversity, such as Madagascar and the Southeast Asia region are recognized for their high tectonic complexity (Pellissier et al. 2018). The study of closely related species distributed across continents serves as a valuable approach for characterizing the evolutionary trajectories of populations. It allows us to discern the impacts of environmental factors and climate change, providing insights into how geographical events have shaped the current biodiversity. The study of such species involves understanding the mechanisms, timing, and contributing factors that led to divergence, ultimately resulting in the formation of distinct taxa-essentially, the process of speciation.

One such example of intercontinental divergence and speciation is the Woolly-necked storks (collective term for Asian Woollynecks and African Woollynecks), which are distributed across Africa including sub-Saharan Africa, and Asia including the southeastern islands (Birdlife International 2023). Their range extends from Senegal and Ethiopia in Africa, to India, Sri Lanka, and Southeast Asia (Fig. 1A) (BirdLife International and Handbook of the Birds of the World 2022). The taxonomic classification of Woolly-necked storks has been a matter of ongoing debate and uncertainty, with limited knowledge about the diversity of these birds across their extensive distribution (Mlodinow et al. 2022). Such ambiguity holds potential implications for the conservation of these species (Sundar 2020). African and Asian Woollynecks were previously regarded as a single species until 2010. However, due to observed distinctions in morphology and plumage coloration, two separate species were officially recognized: the Asian Woollynecks *(Ciconia episcopus)* and the African Woollynecks *(Ciconia microscelis)* (Tobias et al. 2010; del Hoyo and Collar 2014).

**Figure 1:**
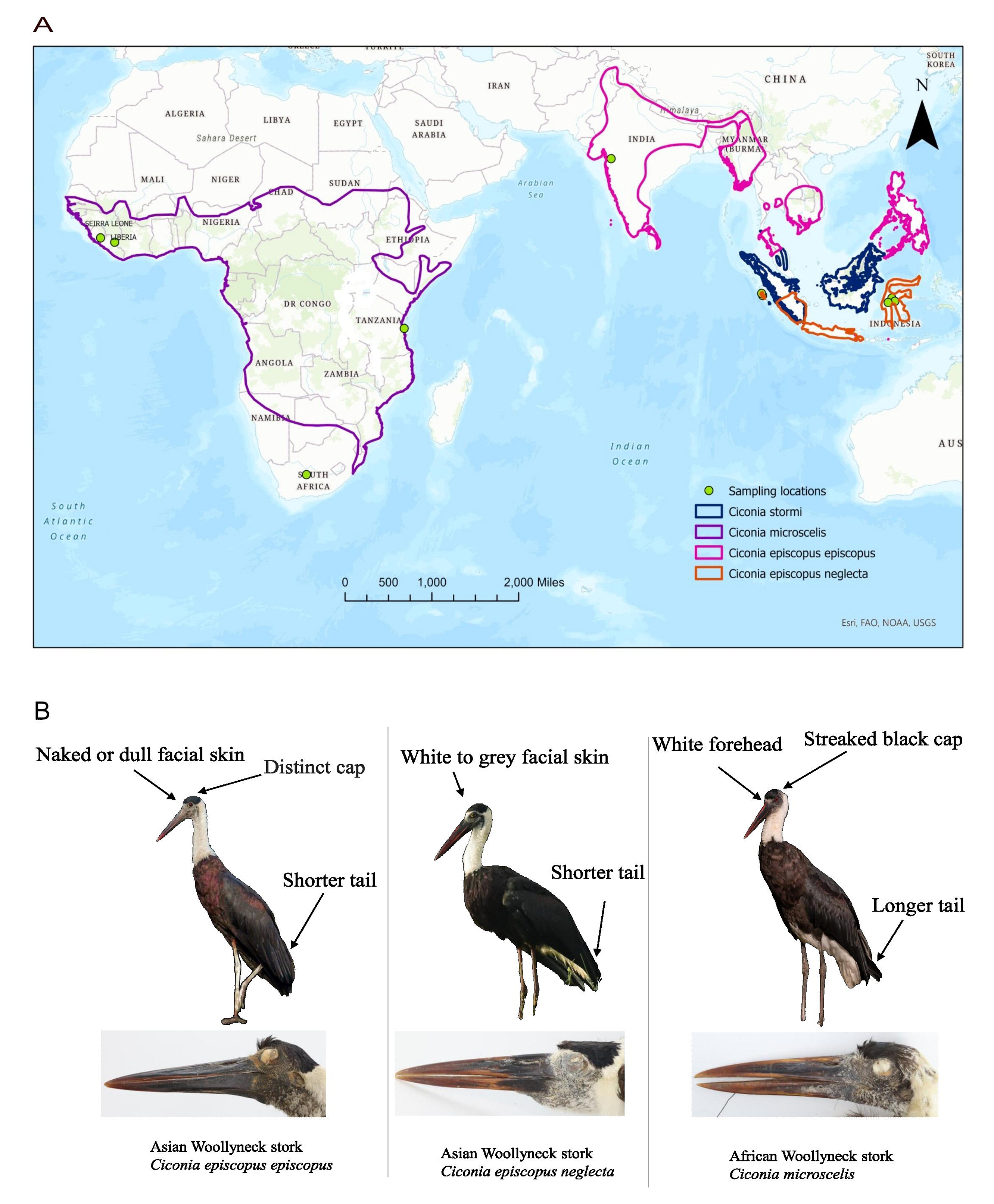
Distribution, sample location, and morphology of storks used in this study: **(A)** distribution of African and Asian Woolly-necked storks and Storm’s storks. Species and subspecies distribution polygons were made using publicly available data from (BirdLife International and Handbook of the Birds of the World 2022). The distribution of *C. e. neglecta* is based on data from Mlodinow et al. 2022. Green dots correspond to the sampling location. **(B)** Morphological differences between Woolly-necked storks. Photographs © Vijayandra Desai, Leonardus Adi Saktyari (ML425347741) and Jonah Gula (left to right, Bird photos) and Jeremiah Trimble, Museum of Comparative Zoology, Harvard University (headshot of museum specimens).

Asian/African Woollynecks are large wading birds, measuring approximately 86 to 95 centimeters in height, with a predominantly black body and a white neck that appears fluffy due to elongated feathers (Birdlife International 2023). Asian Woollynecks have a characteristic distinct black cap with naked or dull facial skin and a shorter tail, while African Woollynecks have a streaked black cap with white facial skin and a longer tail (Mlodinow et al. 2022)(Fig. 1B). While Asian and African Woollynecks are predominantly associated with wetland and farmland habitats, there is an increasing observation of them utilizing urban environments (Thabethe and Downs 2018; Hasan and Ghimire 2020). The urban association is particularly pronounced in the African Woollynecks in Durban, South Africa, where they venture into residential areas and feed on human-provided food (Thabethe and Downs 2018). The re-classification of Woolly-necked storks into two distinct species has had a significant impact on their conservation status. The conservation status of Asian Woollynecks shifted from “Least Concern” to “Vulnerable” in 2014 (Birdlife International 2020). Presently, it is considered “Near Threatened” based on recent population estimations (Kittur and Sundar 2020). In contrast, the conservation status of African Woollynecks has remained unchanged, continuing to be designated as “Least Concern” (Birdlife International 2020).

African Woollynecks appear morphologically homogeneous, and their population size is generally considered to be stable throughout the distribution range (Birdlife International 2020). Contrastingly, Asian Woollynecks are categorized into two sub-species due to discernible morphological differences; *Ciconia episcopus episcopus*, located in South Asia, the Maley Peninsula, and mainland Southeast Asia, and *Ciconia episcopus neglecta*, found in Southern Sumatra, Java, and Wallacea of Indonesia (Fig. 1A) (Gill et al. 2022; Mlodinow et al. 2022). Both subspecies differ in the color of their facial skin and iris (del Hoyo et al. 2020). The demographic status of Asian Woollynecks varies significantly across their distribution range. Populations in India, Sri Lanka, and Nepal have shown relative stability or even an increase (Ghimire et al. 2021) compared to Southeast Asia such as the Philippines, where these birds have faced localized extinctions, potentially linked to habitat deterioration, wetland degradation, pollution, and increased hunting activities (Birdlife International 2020; Kittur and Sundar 2020; Ghimire et al. 2021). Furthermore, the classification of *C. e. neglecta* and *C. e. episcopus* as subspecies is currently under debate, primarily due to the scarcity of reliable information regarding their phenotypic, genetic, and ecological diversity. Additionally, the boundaries of the distribution, including whether there is contact or isolation between each subspecies, and differences in ecology remain unclear, particularly in Southeast Asia (Mlodinow et al. 2022).

Currently, the taxonomic classification of Woolly-necked stork species and subspecies is exclusively based on morphological traits (Mlodinow et al. 2022). This limitation hinders a comprehensive understanding of the transcontinental evolution of these birds and may significantly impact conservation efforts directed toward their populations (Gutiérrez and Helgen 2013; Zachos 2013). Genomic studies of these species hold the potential to not only support their taxonomic status and characterize their evolutionary history but also to offer resources for in-depth exploration of processes underlying their adaptation to local environments throughout their wide distribution range. In this study, we employed whole-genome sequencing in museum specimens to investigate the genetic diversity of Woolly-necked storks and provide insights into their taxonomy and evolutionary history. Our findings contribute to a discussion on their taxonomic status and its potential implications for conservation efforts, population management, and assessments of conservation status.

## Materials and methods

### Obtaining museum specimens

We conducted a search query using the keyword “Ciconia” in the Vertnet online database—a platform for sharing and accessing biodiversity data from diverse biological collections worldwide (http://vertnet.org/). Our search identified specimens of both Asian and African Woollynecks housed at the Museum of Comparative Zoology (MCZ) at Harvard University. From the MCZ, we obtained toe-pad skin samples from a total of 13 museum specimens, including six from Asian Woollynecks *(Ciconia episcopus)*, encompassing both sub-species (three *C. e. episcopus* and three *C. e. neglecta*), five from African Woollynecks *(Ciconia microscelis)*, and two from Storm’s stork *(Ciconia stormi)*, which we utilized as an outgroup. Storm’s (Mlodinow et al. 2022). These specimens were collected from various locations in Asia and Africa between 1934 and 1953 (Supplementary Table 1).

### DNA extraction and whole genome sequencing

DNA extraction from the toe-pad skin samples was performed using the DNeasy Blood and Tissue Kit (Qiagen, Valencia, CA, USA) with a modified version of the protocol suggested by Harvey et al. 2020. Given that the samples were over 70 years old, we adhered to the recommended guidelines for ancient DNA analysis to mitigate the risk of environmental contamination (Orlando et al. 2021). To prevent cross-contamination, all laboratory procedures were conducted in a biosafety cabinet dedicated exclusively to ancient DNA analysis, where no fresh samples were processed. Additionally, all equipment used in the extraction process was carefully cleaned and sterilized before use.

The quantification of extracted DNA was performed using a Qubit fluorometer (Thermo-Fisher, Waltham, MA, USA). However, one sample from the Asian Woolly-necked storks yielded an insufficient amount of DNA and was consequently excluded from subsequent procedures. The DNA from the remaining 12 samples underwent library preparation and whole-genome sequencing, carried out by a commercial genome sequencing provider, Admera Health, New Jersey, United States. The DNA libraries, with an average fragment size of about 400 bp, were sequenced using the Illumina NovaSeq platform, generating 150 bp paired-end reads. The sequencing process aimed to achieve approximately 10X coverage for each sample.

### Sequence alignment and variant calling

All obtained reads were quality-checked using FASTQC (https://www.bioinformatics.babraham.ac.uk/projects/fastqc/). We then mapped the reads from each individual against the reference genome of Maguari stork (*Ciconia maguari),* (Bioproject PRJNA715733) (Feng et al. 2020) using BWA v0.7.17 with default parameters (Li and Durbin 2009). The Maguari stork genome was the closest relative of Woolly-necked storks available in public databases at the time of the analysis. We checked the alignments for PCR duplicates using PICARD (http://picard.sourceforge.net/) and used the Genome Analysis Toolkit (GATK) v4.2.0.0 (McKenna et al. 2010) and GATK best practice recommendations (Van der Auwera et al. 2013) for base quality recalibrations, insertion/deletion (INDEL) realignment, single nucleotide polymorphisms (SNPs) and INDELs discovery, and genotyping across all 12 samples. Analysis of SNP quality parameters was done according to an in-house pipeline previously developed in the lab (Lamichhaney et al. 2015). We excluded INDELs and set up the following parameters in GATK to remove low-quality SNPs: FS > 100.0, MQ < 50.0, MQRankSum < -5.0, ReadPosRankSum < -4.0, BaseQRankSum < -5.0, DP > 500, DP < 10, QD < 5.0, Q > 100 and GQ >10 (see details on each parameter here - https://gatk.broadinstitute.org/hc/en-us/articles/360035531692-VCF-Variant-Call-Format). We further calculated the percentage of missing SNPs across samples using VCFTools v0.1.13 (Danecek et al. 2011) and kept only the SNPs present in all 12 samples for downstream analysis.

### Phylogeny reconstruction and estimates of genetic diversity

We generated an approximate maximum-likelihood phylogeny with FastTree v0.2.1 with recommended default parameters for nucleotide alignments (Price et al. 2010). To assess the local support for each node in the phylogenetic tree, we employed the Shimodaira-Hasegawa test implemented in FastTree. We assessed genetic diversity within and between species with VCFtools v0.1.13 (Danecek et al. 2011) computing genetic metrics such as nucleotide diversity (pi), heterozygosity, inbreeding coefficient (F), and fixation index (F_ST_). For the calculation of nucleotide diversity and inbreeding coefficient (F), we made separate VCF files with SNPs present in at least one sample within that population.

### Population structure and ancestry

We examined the genetic differentiation among samples using a Principal Component Analysis (PCA) with Plink v.1.9 (Chang et al. 2015). As one major assumption of a PCA is independent data, we first pruned our SNP dataset considering linkage disequilibrium (LD). We removed any SNP that shows an r^2^ > 0.1 within a 50 kb window and step size of 10 bp and kept a total of 1780 SNPs for the PCA. We plotted the PCA output in R (Team 2016).

We further estimate the ancestry of individuals based on a genome-wide SNPs dataset using Admixture v.1.3.0 (Alexander et al. 2009) which utilizes a maximum-likelihood approach. To determine the optimal number of genetically distinct clusters (K) that best represent our data, we conducted an exploratory analysis using a cross-validation procedure. This analysis was performed with the K-means method implemented in Admixture. We generated a plot to visualize the proportion of ancestry attributed to each cluster and sample using R (Team 2016).

### Examine the evidence of gene flow between species

We performed two separate analyses to examine the evidence of possible gene flow among Woolly-necked storks. First, we used Treemix v0.1.13 (Pickrell and Pritchard 2012) which utilizes allele frequency information from each SNP to quantify genetic relationships and identify instances of admixture within the branches of the phylogenetic tree. To determine the optimal number of migration events, we performed an analysis using 500 bootstrap replicates of the SNP data. For each of the 1-5 different migration events, we conducted 10 replicates (see results and Supplementary data for details). Second, we utilized ABBA-BABA tests (Green et al. 2010; Durand et al. 2011) and computed Patterson’s D-statistics and f4-ratio statistics in Dsuite (Malinsky et al. 2021) using Storm’s stork as an outgroup.

## Results

### Quality of DNA from historic museum specimens and their resulting genomic data

The amount and quality of DNA obtained, along with the quality of the resulting genome sequence data, are critical aspects in genomic studies involving historic museum specimens. We obtained sufficient DNA for whole genome sequencing using toe-pad skin samples from 12 (out of 13) museum specimens (Supplementary Table 2). The amount of whole genome sequence data generated from these samples was as expected (∼10 Gb of genomic reads per sample, Supplementary Table 2). Various sequence quality parameters such as (a) per base sequence quality (b) per sequence quality scores (c) per base sequence content and (d) per sequence GC content were indicative of good quality sequence data (Supplementary Fig. 1). However, the fragment size of the genomic DNA extracted from all 12 samples was low, indicating DNA degradation (Supplementary Fig. 2), which is expected for DNA obtained from dry toe-pad skin samples collected more than 70 years ago.

Alignment of reads to the reference genome of the Maguari stork (*Ciconia maguari)*, the sequence mapping rate ranged from 83.2% to 99.34 % across our 12 samples (Supplementary Table 3). One specimen of *Ciconia stormi* (MCZ: 170531) from 1934, the oldest among the samples we collected, had the lowest mapping rate (83.2%). The average genome-wide sequencing depth ranged from 4.5 to 6.9 (Supplementary Table 3). Following our pipeline, we obtained ∼13.5 million good-quality SNPs (see methods for details). However, as the genomic DNA was degraded for most of the specimens (Supplementary Fig. 2), the rate of missing SNPs across samples was high (Supplementary Table 4). We found the number of missing SNPs in a sample was strongly correlated with the quality of the genomic DNA (Supplementary Fig. 3). Hence, to remove bias due to missing SNPs, we followed a conservative approach and only used 33,087 SNPs that were genotyped in all 12 samples (missingness = 0) for downstream analyses.

### Genome-based phylogeny

We constructed a phylogeny using the filtered dataset of 33,087 SNPs, employing a maximum-likelihood approach (Price et al. 2010) and using the Storm’s stork samples to root the tree as an outgroup. The African and Asian woollyneck samples formed two reciprocally monophyletic clades (Fig. 2). Within the Asian clade the samples of each subspecies of Asian Woollyneck, *C. e. neglecta* and *C. e. episcopus* clustered together, demonstrating distinct genetic groups. In the African clade, the branch lengths are shorter, and the node supports are slightly lower than in the Asian clade. African samples also formed two distinct clades but unrelated with the geographical distance among them; Clade 1-samples from South Africa, Sierra-Leone, and Liberia, and Clade 2 – samples from Tanzania clustered together with the sample of unknown origin (Fig. 2, Supplementary Table 1).

**Figure 2:**
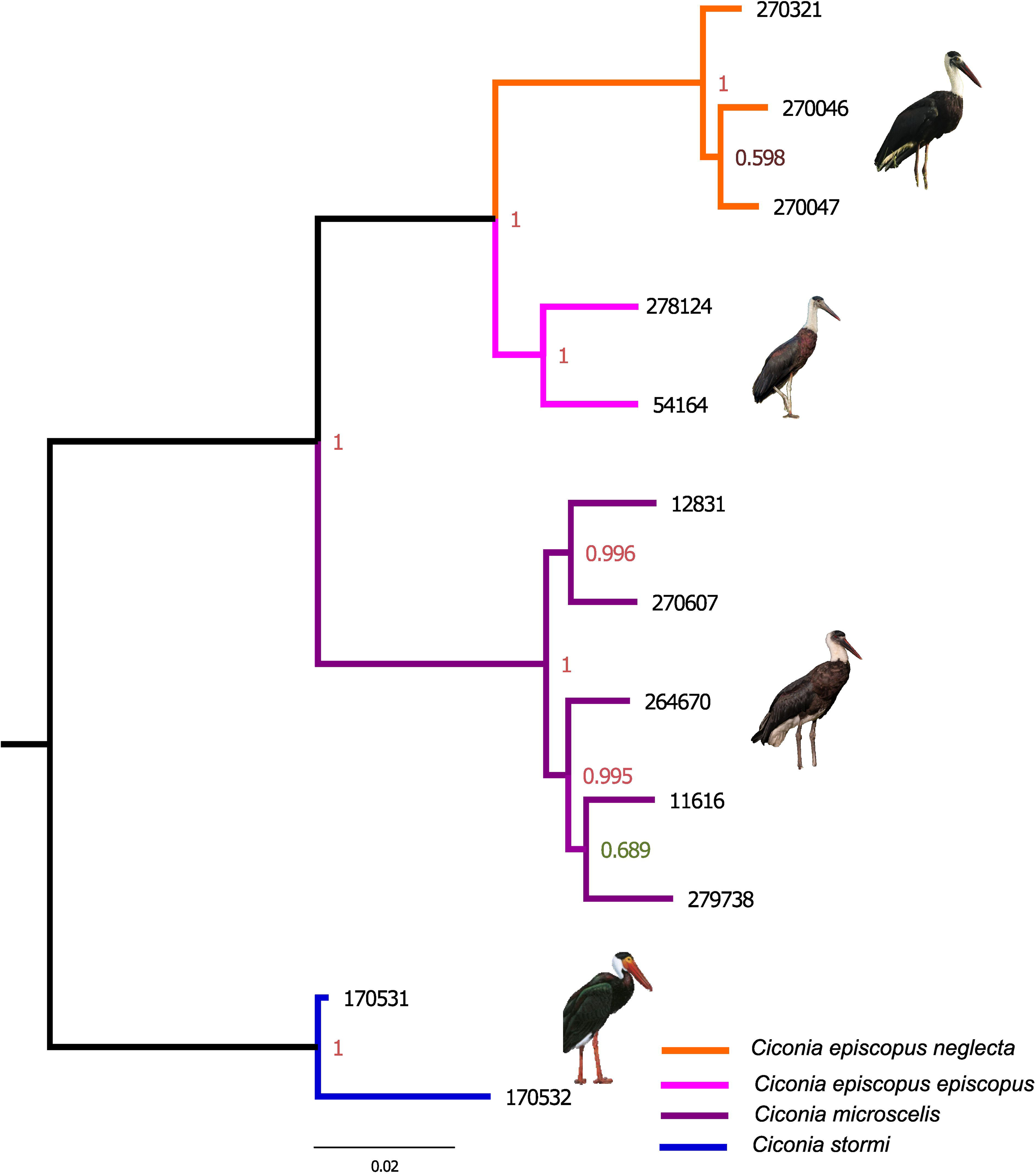
Phylogeny of Woolly-necked storks: Maximum-likelihood phylogenetic tree based on 33,087 genome-wide SNPs. Local support values at each node were calculated based on Shimodaira–Hasegawa test (Price et al. 2010). Photographs © Leonardus Adi Saktyari(ML425347741), Vijayandra Desai, Jonah Gula Francesc Jutglar (Birds of the World) (From top to bottom).

### Genetic diversity among groups

Average genome-wide nucleotide diversity (pi) was lower in the African samples compared to others including the outgroup (Table 1). The inbreeding coefficient (F) was close to zero for all samples (excluding the outgroups) indicating a more outbred population with a broader genetic pool. The average genome-wide genetic divergence (F_ST_) between Asian and African Woolly-necked storks was estimated as 0.20 (Table 2). The F_ST_ between the two Asian subspecies was 0.14 whereas between the two African clades identified from the phylogenetic tree was -0.006. The F_ST_ for Asian and African Woolly-necked stork clades compared with outgroup Storm’s stork were very similar (0.25 and 0.27 respectively). The F_ST_ between *C. e. neglecta* and African samples was higher (0.27) compared to the one between *C. e. episcopus* and African samples (0.16). We also calculated F_ST_ in 15 kb windows across the genome comparing Asian and African Woollynecks and identified 72 genomic windows with high F_ST_ (Fst > 0.8) (Fig. 3).

**Table 1:**
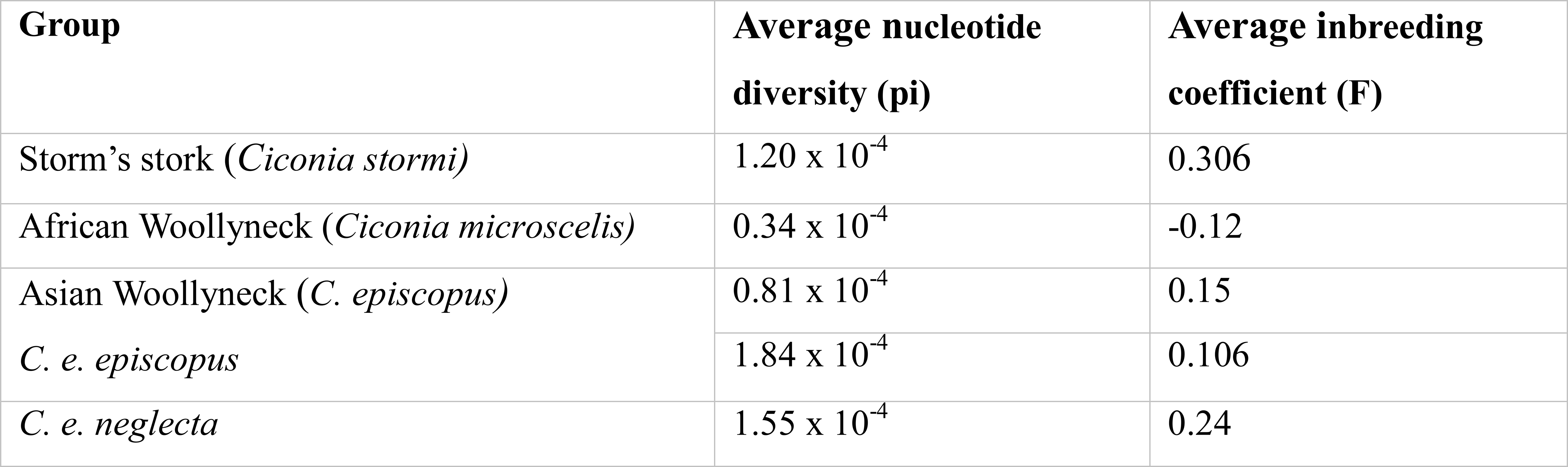
Estimates of genetic diversity.

**Table 2:**
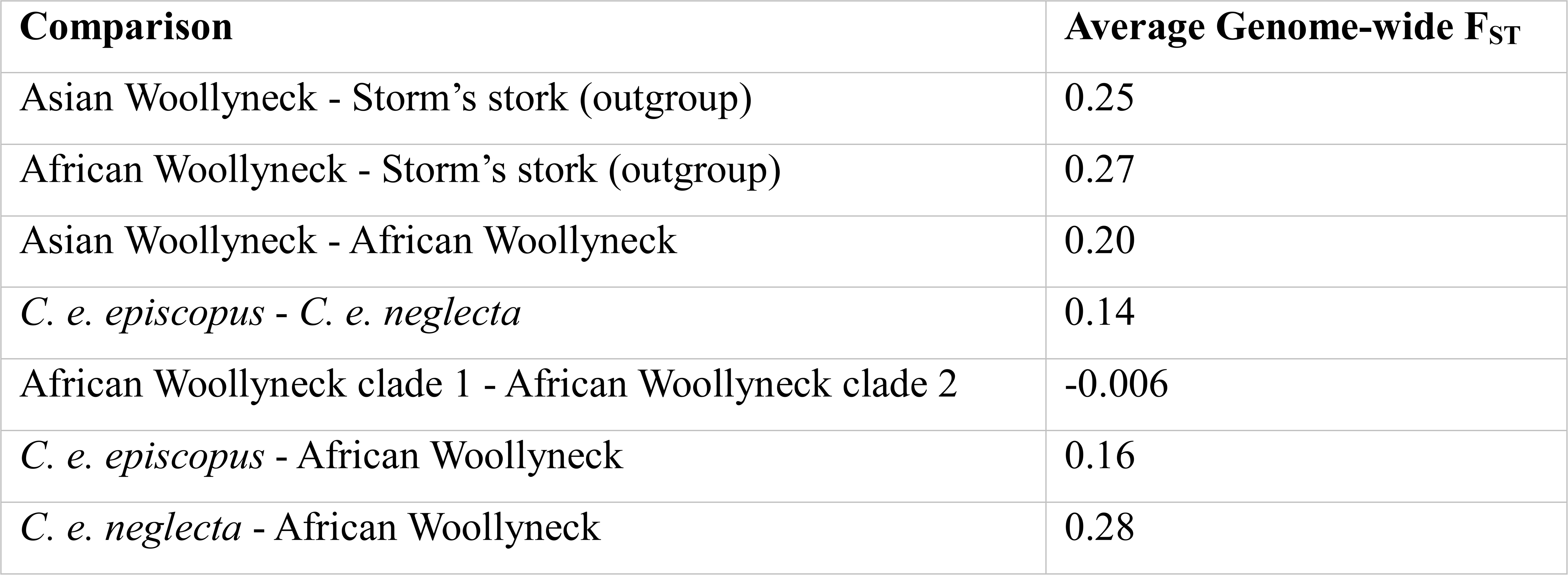
Estimates of average genome-wide genetic divergence (F_ST_).

**Figure 3:**
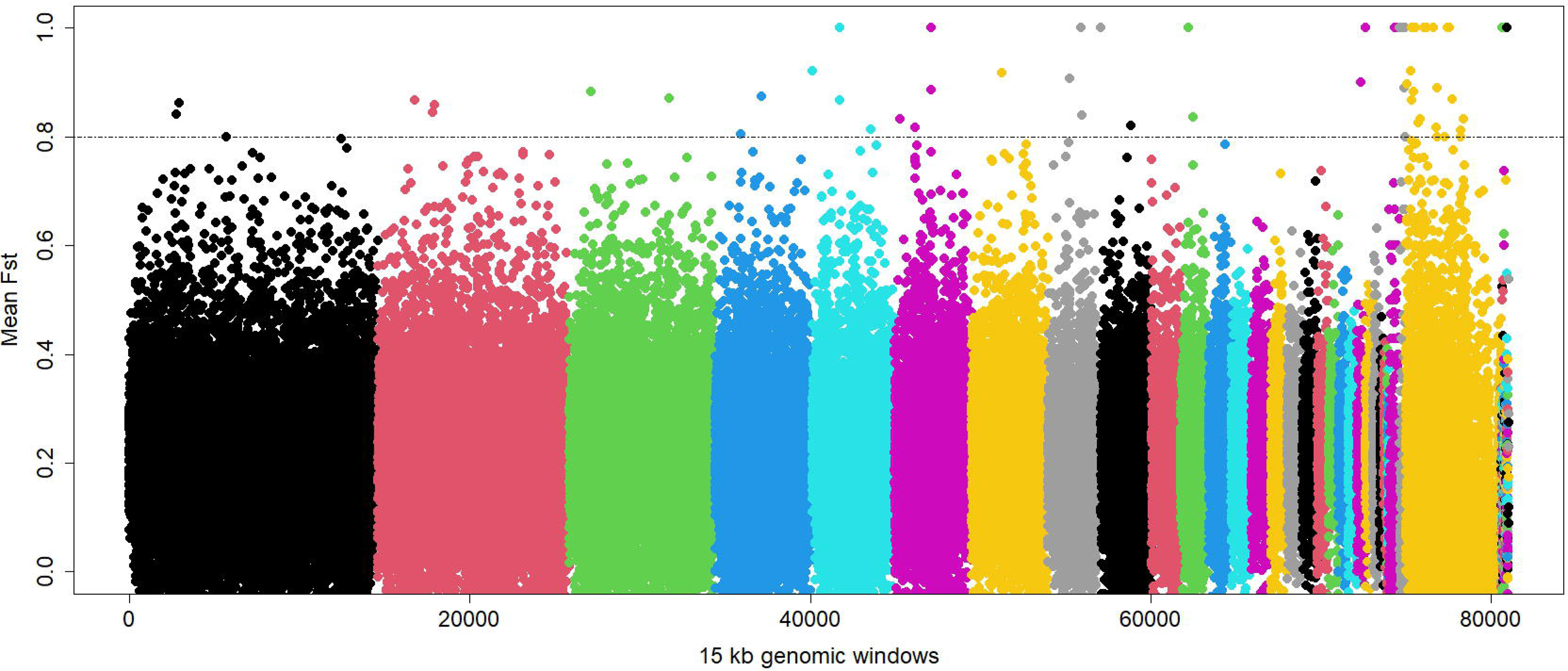
Genomic regions of differentiation between African and African Woollynecks: Genome-wide F_ST_ calculated for each 15 kb genomic window between African and African Woolly-necked storks. All SNPs from a respective chromosome are coded differently. Genomic regions with high F_ST_ (> 0.8) are shown above a dashed horizontal line.

### Population structure and ancestry

Principal component analysis (PCA) indicated first and second principal components (PCs) explained almost 50 % of the genetic variation among the samples (PC1=31.2 % and PC2=15.5 %, Supplementary Fig. 4). The first principal component separates the Asian Woollynecks, the African Woollynecks, and the Storm’s Stork groups, while the second principal component mainly separates the outgroup Storm’s Stork from all Woolly-necked storks (Fig. 4A). Across the first principal component the Asian samples make two distinct genetic clusters corresponding to *C. e. neglecta* and *C. e. episcopus* while the African samples are not differentiated forming a single cluster. In a PCA excluding the outgroup Storm’s Stork, the differences between *C. e. neglecta* and *C. e. episcopus* are even more evident while the African samples still make a single tight cluster (Fig. 4B).

**Figure 4:**
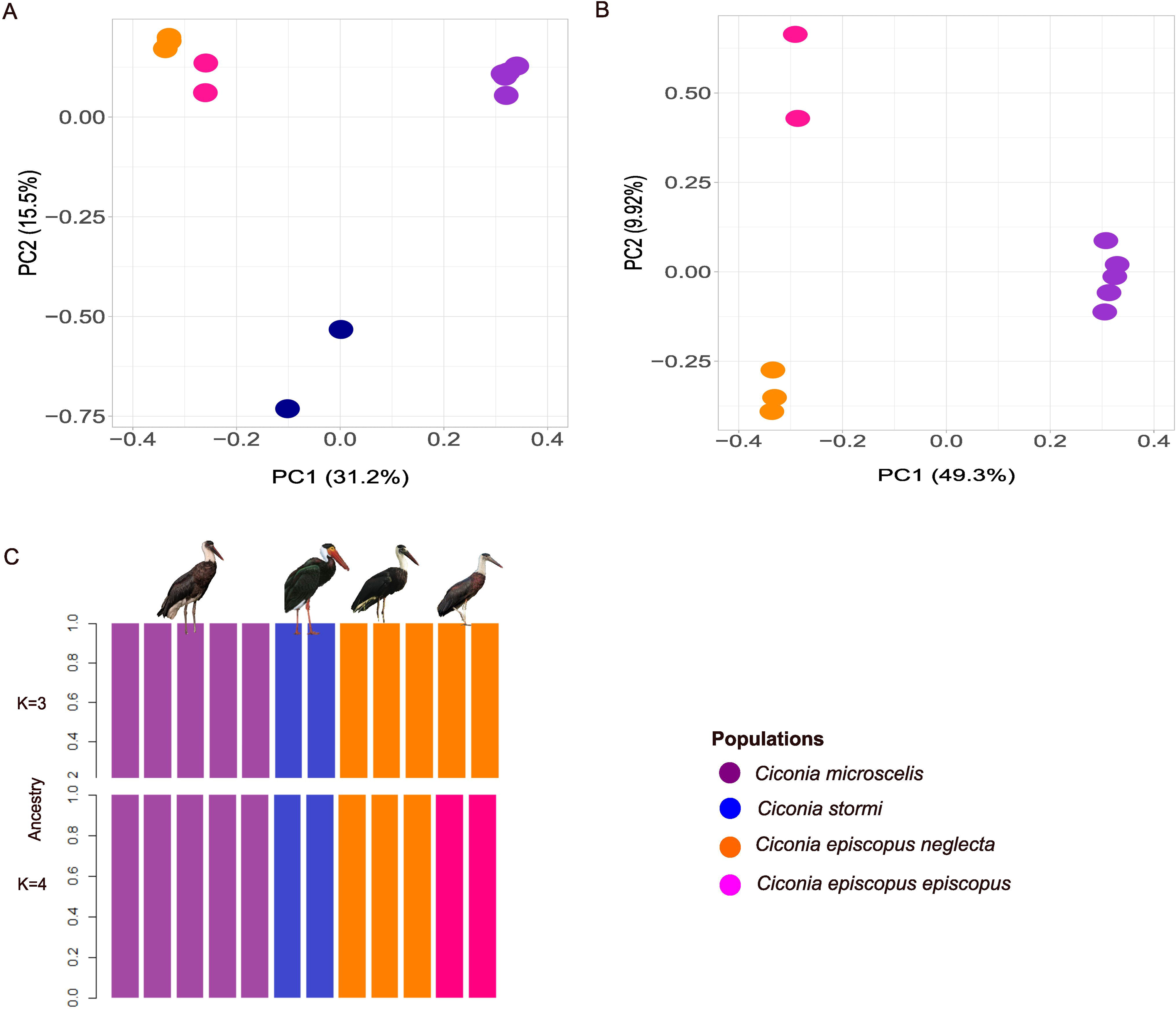
The population structure of Woolly-necked storks was assessed through Principal Component Analysis (PCA), both with **(A)** and without **(B)** the outgroup Storm’s stork. The results reveal distinct clusters for African Woollynecks, Asian Woollynecks, and Storm’s stork, and differentiate between the Asian subspecies *C. e. neglecta* and *C. e. episcopus*. **(C)** The Admixture plot for K=3 illustrates distinct clusters for samples corresponding to Storm’s stork, African Woolly-necked storks, and Asian Woolly-necked storks. Furthermore, for K=4, the plot reveals a finer separation within the Asian Woolly-necked storks, distinguishing between subspecies *C. e. neglecta* and *C. e. episcopus*. Photographs © Jonah Gula, Francesc Jutglar (Birds of the World), Leonardus Adi Saktyari (ML425347741), and Vijayandra Desai, (From left to right).

We assessed various values of K using a cross-validation procedure and identified that K=3 or 4 is optimal for admixture analysis (Supplementary Table 5). K=3 admixture run grouped samples into Asian Woollynecks, African Woollynecks, and the Storm’s stork, whereas the K=4 run further divided the Asian samples into the two groups, consistent with their subspecies status, *C. e. episcopus* and *C. e. neglecta* (Fig. 4C).

### Gene flow among populations

We employed TreeMix (Pickrell and Pritchard 2012) to identify possible evidence of gene flow among populations. A bootstrapping procedure identified migration events of 1 or 2 was optimal for our data (Supplementary Fig. 5). However, the TreeMix analysis did not provide any solid evidence of possible gene flow between populations (Supplementary Fig. 6). To further explore possible evidence of gene flow between the Woolly-necked storks clades we carried out ABBA-BABA analysis to estimate D and f4-ratio statistics in Dsuite (Malinsky et al. 2021) and using Storm’s stork as an outgroup. We did not find evidence of gene flow between Asian Woollyneck *C. episcopus* and *C. microscelis*, but we found evidence of gene flow between the two clades of African Woolly-necked storks (Z score = 9.24, p = 0.00).

## Discussion

In this study, we have conducted genomic analysis on museum specimens collected ∼70 years ago to investigate the evolutionary relationships of Woolly-necked storks, whose taxonomic status has been a topic of uncertainty (Sundar 2020). To our knowledge, this is the first study to produce genomic resources for the Woolly-necked storks and employ the genomic data to examine the existing taxonomy, which has solely relied on morphological data to date. Based on a genome-wide phylogeny, principal component analysis, ancestry analysis, and various metrics of genetic diversity, our results have revealed that Asian and African Woollynecks are genetically distinct sister clades, lending support for the current taxonomy of these birds which recognizes Asian and African Woollynecks as distinct species, *Ciconia episcopus* and *Ciconia microscelis* (del Hoyo et al. 2020; Mlodinow et al. 2022).

Our results also confirmed that Asian Woollynecks are composed of two genetically distinct clades corresponding to the currently recognized subspecies classification, *C. episcopus neglecta,* and *C. episcopus episcopus*. These two groups were differentiated in all the analyses we performed: they formed two distinct clusters in the phylogenetic tree, grouped independently of each other in the principal component analyses, showed high genetic diversity, and were recognized as different groups in admixture analyses. This evidence, in addition to their known morphologic differences and distribution(Mlodinow et al. 2022), suggests *C. e. neglecta* and *C. e. episcopus* are in the process of being independent lineages and may have accumulated enough differences to be recognized as different species, especially considering we did not find evidence of gene flow between them. This process of genetic divergence in these two clades of Asian Woollynecks is possibly boosted by the geographical isolation of *C. e. neglecta* in the southeast islands of Asia in association with the tectonic activity in this zone during the last 10 million years (Verstappen 1997; Hall 2009). While it is crucial to approach the interpretation of our results with caution due to the limited sample size, it is intriguing to note that, with comparable sample sizes, the genetic divergence between *C. e. neglecta* and *C. e. episcopus* is notably higher than the variation observed among African Woollyneck populations, which were collected across the continent. However, comprehensive population genomics studies using additional populations from other Southeast Asian areas, such as other parts of Indonesia, Thailand, Malaysia, and Cambodia will be necessary to evaluate whether these two clades on Asian Woollyneck *C. e. neglecta* should be continued to be classified as subspecies or potentially regarded as distinct species. Analyzing the genetic characteristics of these populations could also offer valuable insights into the substantial ongoing decline of Woolly-necked storks in Southeast Asia (Ghimire et al. 2021).

African Woollyneck populations demonstrated lower genetic diversity in comparison to Asian Woollynecks, as evidenced by a low genome-wide average nucleotide diversity, shorter branches in the phylogeny, and an absence of distinct structure in admixture analysis. These outcomes may suggest that the effective population size of African Woollyneck was smaller during the period when these museum samples were collected, around the 1900s. Alternatively, these results could imply ongoing gene flow among African Woollyneck populations after a recent period of divergence. Due to the lack of reliable data on population size changes in African Woollyneck over the past century, and the limited availability of robust population genomics data, we are unable to provide a more specific explanation. However, anecdotal evidence indicates that African woollynecks were regarded as endangered in South Africa until the 1990s because of the reduced population size (Clancey 1964; Cyrus and Robson 1980), which is consistent with our results.

While we did not detect evidence of population structure among African samples, our phylogenetic analysis revealed that they form two well-supported clades. Contrary to the expectation that populations in closer geographical proximity would exhibit greater genetic similarity, our findings indicated that individuals from Sierra Leone are closely related to those from South Africa, despite the substantial geographical distance between these two regions. A recent study on African Woollyneck populations has suggested that their distribution is fragmented (Gula et al. 2020), particularly in West Africa. The observed phylogenetic pattern in African Woollyneck could potentially be attributed to this fragmented distribution.

The current taxonomy recognizing African Woollyneck and Asian Woollyneck as independent lineages is supported not only by morphological and ecological differences but also by genome-wide differentiation. They were distinct in all our analyses, and we found no evidence of gene flow between them. Despite Wolly-necked storks can fly long distances, it is possible that they do not fly extensively across open water, suggesting water bodies may have played a significant role in the divergence of Asian and African species and the differentiation and likely divergence between *C. e. neglecta* and *C. e. episcopus*. The lack of gene flow may have contributed to reduced genetic variability and decreased population size, for example in the Philippines, where the Asian Woollyneck seems to have been locally extinct from Luzon and neighboring islands (Ghimire et al. 2021).

In a broader context, the phylogenetic relationships among various stork species remain unclear due to the absence of a comprehensive phylogenetic reconstruction that encompasses all species and subspecies within the *Ciconia* genus. Recently a phylogeny based on mitochondrial markers (*COI* and *CytB*) and cytogenetic data placed Storm’s storks as the sister clade of *C. episcopus* (de Sousa et al. 2023) but the African Woollynecks were not included in this phylogeny.

Furthermore, limited information is available regarding the time of divergence among species within the *Ciconidae* family. Although the genus is estimated to be around 9 million years old (http://timetree.org/), the specific divergence times for many species remain largely unknown. These timeframes are notably more recent than the estimated collision between the African plate and the Arabia-Eurasia plates, which is estimated to have occurred between 35 and 25 million years ago (McQuarrie and van Hinsbergen 2013). A general expectation is the divergence between African and Asian Woolly-necked storks should have started when the two continents were already connected. Numerous examples across various taxa, including plants, animals, and microorganisms, illustrate continental-scale divergences, such as the case of wild elephants in Asia and Africa (Fernando et al. 2000; Eggert et al. 2002) and fungi across North America and Europe (Tremble et al. 2023). The study of these widely distributed species contributes to our understanding of the mechanistic basis of divergence and speciation shaped by the Earth’s geological processes across continents (Descombes et al. 2017).

Multiple lines of evidence suggest that both Asian and African Woollynecks are responding to changes in local environments and potentially undergoing local adaptation. Firstly, they demonstrate variations in response to climatic variables, including bioclimatic factors like precipitation and temperature (Gula et al. 2020). For instance, the probability of the presence of Asian Woollynecks increases with precipitation in the warmest quarter, whereas the trend is the opposite for African Woollynecks. The same study predicts a positive association of African Woollynecks with flooded forest/shrubland and grassland with woody vegetation, while Asian Woollynecks are associated with natural vegetation patches and croplands.

Secondly, there is evidence indicating that both species are utilizing urban environments. African Woollynecks have been observed using supplementary feeding in urban areas of South Africa (Thabethe and Downs 2018), and Asian Woollynecks have been documented utilizing artificial structures for nesting in recent years (Vaghela et al. 2015; Hasan and Ghimire 2020). However, African Woollynecks seem to fare better in urban environments than Asian Woollynecks. Given that our samples date back approximately 70 years and our sample size is limited, our ability to draw definitive conclusions about local adaptation is constrained. Nevertheless, we have identified numerous genomic regions showing strong genetic divergence between Asian and African Woollynecks that may potentially harbor genes involved in local adaptation.

This study is anticipated to create avenues for future genomic research that use a larger sample size from populations across the distribution of Asian and African Woollynecks and examine genomic signatures of local adaptation associated with changes in their morphology, the timing of reproduction, and urbanization. We are also suggesting the need to study the diversity of Asian Woollyneck populations to determine if *C. e. neclecta* and *C. e. episcopus* are independent lineages, i.e., species, and therefore they require targeted conservation strategies to mitigate the decline and promote the recovery of Woolly-necked storks in Southeast Asia. Hence, upcoming research endeavors should concentrate on identifying recent indications of habitat overlap or genetic interchange between these two groups. While this study presents evidence of genomic differentiation between Asian and African Woollynecks, the specific evolutionary processes that led to their divergence and the molecular mechanisms underlying their speciation remain topics for future investigation. Studies on patterns of genetic differentiation, phylogenetics, and ecological adaptations in closely related species across continents allow us to better understand how geological events, geographical barriers, and environmental differences shape biodiversity. Woolly-necked Storks are a suitable system to explore these evolutionary dynamics.

## Author Contributions

S.L. and P.G. conceived the study. J.T. contributed to getting access to toe-pad skin samples and took high-resolution photographs of these museum specimens. P.G. performed the laboratory work and bioinformatics analysis with contributions from C.P. P.G., C.P., and S.L. wrote the paper with input from all authors. All authors approved the manuscript before submission.

## Acknowledgments

We thank Dr. Breda Zimkus (Director of collection operations at the Museum of Comparative Zoology at Harvard University) for her assistance in collecting toe-pad skin samples from museum specimens. We thank Teresa Raba, an undergraduate student at Kent State University for her assistance with DNA extractions from museum specimens. We also thank Prof. Prof. Sandi Willows-Munro (University of KwaZulu-Natal, South Africa) for her helpful comments and suggestions on the manuscript. We thank Vijayandra Desai, Jonah Gula, Francesc Jutglar and Leonardus Adi Saktyari, Jeremiah Trimble, and Macauley Library at Cornell Lab of Ornithology for the permit to use their photographs in this paper. We are thankful to Birdlife International for providing distribution-related spatial data on storks. The project was supported by the Department of Biological Sciences, at Kent State University. The authors declare no conflicts of interest.

## Data and script availability

All short-read whole-genome sequencing data have been submitted to NCBI under BioProject Accession number PRJNA996906. Scripts and workflow for all analyses done in this paper will be available on our dedicated GitHub page (https://github.com/sangeet2019/Woollyneck-storks) after the publication of the paper.

## References

Abbott, R., D. Albach, S. Ansell, J. W. Arntzen, S. J. E. Baird, N. Bierne, J. Boughman, A. Brelsford, C. A. Buerkle, R. Buggs, R. K. Butlin, U. Dieckmann, F. Eroukhmanoff, A. Grill, S. H. Cahan, J. S. Hermansen, G. Hewitt, A. G. Hudson, C. Jiggins, J. Jones, B. Keller, T. Marczewski, J. Mallet, P. Martinez-Rodriguez, M. Möst, S. Mullen, R. Nichols, A. W. Nolte, C. Parisod, K. Pfennig, A. M. Rice, M. G. Ritchie, B. Seifert, C. M. Smadja, R. Stelkens, J. M. Szymura, R. Väinölä, J. B. W. Wolf, and D. Zinner. 2013. Hybridization and speciation. J. Evol. Biol. 26:229–246. John Wiley & Sons, Ltd (10.1111).

Alexander, D. H., J. Novembre, and K. Lange. 2009. Fast model-based estimation of ancestry in unrelated individuals. Genome Res. 19:1655–1664. Cold Spring Harbor Laboratory Press.

Bickford, D., D. J. Lohman, N. S. Sodhi, P. K. Ng, R. Meier, K. Winker, K. K. Ingram, and I. Das. 2007. Cryptic species as a window on diversity and conservation. Trends Ecol. Evol. 22:148–155. Elsevier.

Birdlife International. 2020. Ciconia episcopus (Asian Woollyneck). Birdlife International. 2023. The IUCN Red List of Threatened Species.

BirdLife International, and Handbook of the Birds of the World. 2022. Bird species distribution maps of the world. Version 2022.1.

Bolnick, D. I., A. K. Hund, P. Nosil, F. Peng, M. Ravinet, S. Stankowski, S. Subramanian, J. B. Wolf, and R. Yukilevich. 2023. A multivariate view of the speciation continuum. Evolution 77:318–328. Oxford University Press US.

Bonaccorso, E., I. Koch, and A. T. Peterson. 2006. Pleistocene fragmentation of Amazon species’ ranges. Divers. Distrib. 12:157–164.

Campbell, C. R., J. W. Poelstra, and A. D. Yoder. 2018. What is Speciation Genomics? The roles of ecology, gene flow, and genomic architecture in the formation of species. Biol. J. Linn. Soc. 124:561–583.

Card, D. C., B. Shapiro, G. Giribet, C. Moritz, and S. V. Edwards. 2021. Museum Genomics. Annu. Rev. Genet. 55:633–659.

Carney, J. P., and T. A. Dick. 2000. The historical ecology of yellow perch (Perca flavescens[Mitchill]) and their parasites. J. Biogeogr. 27:1337–1347.

Chang, C. C., C. C. Chow, L. C. Tellier, S. Vattikuti, S. M. Purcell, and J. J. Lee. 2015. Second-generation PLINK: rising to the challenge of larger and richer datasets. GigaScience 4:7.

Clancey, P. 1964. The birds of Natal and Zululand. Oliver and Boyd, Edinburgh.

Coyne, J. A. 1992. Genetics and speciation. Nature 355:511–515. Nature Publishing Group UK London.

Craw, D., P. Upton, C. P. Burridge, G. P. Wallis, and J. M. Waters. 2016. Rapid biological speciation driven by tectonic evolution in New Zealand. Nat. Geosci. 9:140–144. Nature Publishing Group.

Cyrus, D., and N. Robson. 1980. Bird Atlas of Natal. University of Natal Press, Pietermaritzburg.

Danecek, P., A. Auton, G. Abecasis, C. A. Albers, E. Banks, M. A. DePristo, R. E. Handsaker, G. Lunter, G. T. Marth, S. T. Sherry, G. McVean, R. Durbin, and 1000 Genomes Project Analysis Group. 2011. The variant call format and VCFtools. Bioinformatics 27:2156–2158.

Darwin, C. 1859. On the Origin of Species by Means of Natural Selection, Or, The Preservation of Favoured Races in the Struggle for Life. J. Murray.

De Queiroz, K. 2007. Species Concepts and Species Delimitation. Syst. Biol. 56:879–886.

de Sousa, R. P. C., P. S. B. Campos, M. da S. dos Santos, P. C. O’Brien, M. A. Ferguson-Smith, and E. H. C. de Oliveira. 2023. Cytotaxonomy and Molecular Analyses of Mycteria americana (Ciconiidae: Ciconiiformes): Insights on Stork Phylogeny. Genes 14:816. Multidisciplinary Digital Publishing Institute.

del Hoyo, J., and N. J. Collar. 2014. HBW and BirdLife International Illustrated Checklist of the Birds of the World. Volume 1. Non-passerines. Lynx Edicions, Barcelona.

del Hoyo, J., A. Elliott, N. Collar, E. Garcia, P. F. D. Boesman, and G. M. Kirwan. 2020. Woolly-necked Stork (Ciconia episcopus), version 1.0. Birds World, doi: 10.2173/bow.wonsto1.01. Cornell Lab of Ornithology, Ithaca, NY, USA.

Descombes, P., F. Leprieur, C. Albouy, C. Heine, and L. Pellissier. 2017. Spatial imprints of plate tectonics on extant richness of terrestrial vertebrates. J. Biogeogr. 44:1185–1197.

Dirzo, R., and P. H. Raven. 2003. Global State of Biodiversity and Loss. Annu. Rev. Environ. Resour. 28:137–167.

Durand, E. Y., N. Patterson, D. Reich, and M. Slatkin. 2011. Testing for ancient admixture between closely related populations. Mol. Biol. Evol., doi: 10.1093/molbev/msr048.

Eggert, L. S., C. A. Rasner, and D. S. Woodruff. 2002. The evolution and phylogeography of the African elephant inferred from mitochondrial DNA sequence and nuclear microsatellite markers. Proc. Biol. Sci. 269:1993–2006.

Engelbrecht, H. M., W. R. Branch, E. Greenbaum, M. Burger, W. Conradie, and K. A. Tolley. 2020. African Herald snakes, Crotaphopeltis, show population structure for a widespread generalist but deep genetic divergence for forest specialists. J. Zool. Syst. Evol. Res. 58:1220–1233.

Felsenstein, J. 1988. Phylogenies from Molecular Sequences: Inference and Reliability. Annu. Rev. Genet. 22:521–565.

Feng, S., J. Stiller, Y. Deng, J. Armstrong, Q. Fang, A. H. Reeve, D. Xie, G. Chen, C. Guo, B. C. Faircloth, B. Petersen, Z. Wang, Q. Zhou, M. Diekhans, W. Chen, S. Andreu-Sánchez, A. Margaryan, J. T. Howard, C. Parent, G. Pacheco, M.-H. S. Sinding, L. Puetz, E. Cavill, Â. M. Ribeiro, L. Eckhart, J. Fjeldså, P. A. Hosner, R. T. Brumfield, L. Christidis, M. F. Bertelsen, T. Sicheritz-Ponten, D. T. Tietze, B. C. Robertson, G. Song, G. Borgia, S. Claramunt, I. J. Lovette, S. J. Cowen, P. Njoroge, J. P. Dumbacher, O. A. Ryder, J. Fuchs, M. Bunce, D. W. Burt, J. Cracraft, G. Meng, S. J. Hackett, P. G. Ryan, K. A. Jønsson, I. G. Jamieson, R. R. da Fonseca, E. L. Braun, P. Houde, S. Mirarab, A. Suh, B. Hansson, S. Ponnikas, H. Sigeman, M. Stervander, P. B. Frandsen, H. van der Zwan, R. van der Sluis, C. Visser, C. N. Balakrishnan, A. G. Clark, J. W. Fitzpatrick, R. Bowman, N. Chen, A. Cloutier, T. B. Sackton, S. V. Edwards, D. J. Foote, S. B. Shakya, F. H. Sheldon, A. Vignal, A. E. R. Soares, B. Shapiro, J. González-Solís, J. Ferrer-Obiol, J. Rozas, M. Riutort, A. Tigano, V. Friesen, L. Dalén, A. O. Urrutia, T. Székely, Y. Liu, M. G. Campana, A. Corvelo, R. C. Fleischer, K. M. Rutherford, N. J. Gemmell, N. Dussex, H. Mouritsen, N. Thiele, K. Delmore, M. Liedvogel, A. Franke, M. P. Hoeppner, O. Krone, A. M. Fudickar, B. Milá, E. D. Ketterson, A. E. Fidler, G. Friis, Á. M. Parody-Merino, P. F. Battley, M. P. Cox, N. C. B. Lima, F. Prosdocimi, T. L. Parchman, B. A. Schlinger, B. A. Loiselle, J. G. Blake, H. C. Lim, L. B. Day, M. J. Fuxjager, M. W. Baldwin, M. J. Braun, M. Wirthlin, R. B. Dikow, T. B. Ryder, G. Camenisch, L. F. Keller, J. M. DaCosta, M. E. Hauber, M. I. M. Louder, C. C. Witt, J. A. McGuire, J. Mudge, L. C. Megna, M. D. Carling, B. Wang, S. A. Taylor, G. Del-Rio, A. Aleixo, A. T. R. Vasconcelos, C. V. Mello, J. T. Weir, D. Haussler, Q. Li, H. Yang, J. Wang, F. Lei, C. Rahbek, M. T. P. Gilbert, G. R. Graves, E. D. Jarvis, B. Paten, and G. Zhang. 2020. Dense sampling of bird diversity increases power of comparative genomics. Nature 587:252–257.

Fernando, P., M. E. Pfrender, S. E. Encalada, and R. Lande. 2000. Mitochondrial DNA variation, phylogeography and population structure of the Asian elephant. Heredity 84:362–372.

Forsman, A. 2015. Rethinking phenotypic plasticity and its consequences for individuals, populations and species. Heredity 115:276–284. Nature Publishing Group.

Gagnaire, P.-A. 2020. Comparative genomics approach to evolutionary process connectivity. Evol. Appl. 13:1320–1334. Wiley Online Library.

Genereux, D. P., A. Serres, J. Armstrong, J. Johnson, V. D. Marinescu, E. Murén, D. Juan, G. Bejerano, N. R. Casewell, L. G. Chemnick, J. Damas, F. Di Palma, M. Diekhans, I. T. Fiddes, M. Garber, V. N. Gladyshev, L. Goodman, W. Haerty, M. L. Houck, R. Hubley, T. Kivioja, K.-P. Koepfli, L. F. K. Kuderna, E. S. Lander, J. R. S. Meadows, W. J. Murphy, W. Nash, H. J. Noh, M. Nweeia, A. R. Pfenning, K. S. Pollard, D. A. Ray, B. Shapiro, A. F. A. Smit, M. S. Springer, C. C. Steiner, R. Swofford, J. Taipale, E. C. Teeling, J. Turner-Maier, J. Alfoldi, B. Birren, O. A. Ryder, H. A. Lewin, B. Paten, T. Marques-Bonet, K. Lindblad-Toh, E. K. Karlsson, and Zoonomia Consortium. 2020. A comparative genomics multitool for scientific discovery and conservation. Nature 587:240–245.

Ghimire, P., R. Ghimire, M. Low, B. S. Bist, and N. Pandey. 2021. The Asian Woollyneck Ciconia episcopus: A review of its status, distribution and ecology. Ornithol. Sci. 20:223–233.

Gill, F. B., D. Donsker, and P. Rasmussen. 2022. Storks, frigatebirds, boobies, darters, cormorants – IOC World Bird List.

Goodwin, S., J. D. McPherson, and W. R. McCombie. 2016. Coming of age: ten years of next-generation sequencing technologies. Nat. Rev. Genet. 17:333–351. Nature Publishing Group.

Grant, P. R. G. B. R. 2014. 40 Years of Evolution: Darwin’s Finches on Daphne Major Island. Princeton University Press.

Green, R. E., J. Krause, A. W. Briggs, T. Maricic, U. Stenzel, M. Kircher, N. Patterson, H. Li, W. Zhai, M. H.-Y. Fritz, N. F. Hansen, E. Y. Durand, A.-S. Malaspinas, J. D. Jensen, T. Marques-Bonet, C. Alkan, K. Prüfer, M. Meyer, H. A. Burbano, J. M. Good, R. Schultz, A. Aximu-Petri, A. Butthof, B. Höber, B. Höffner, M. Siegemund, A. Weihmann, C. Nusbaum, E. S. Lander, C. Russ, N. Novod, J. Affourtit, M. Egholm, C. Verna, P. Rudan, D. Brajkovic, Ž. Kucan, I. Gušic, V. B. Doronichev, L. V. Golovanova, C. Lalueza-Fox, M. de la Rasilla, J. Fortea, A. Rosas, R. W. Schmitz, P. L. F. Johnson, E. E. Eichler, D. Falush, E. Birney, J. C. Mullikin, M. Slatkin, R. Nielsen, J. Kelso, M. Lachmann, D. Reich, and S. Pääbo. 2010. A draft sequence of the Neandertal genome. Science 328:710–722.

Gula, J., K. S. G. Sundar, and W. R. J. Dean. 2020. Known and potential distributions of the African Ciconia miscroscelis and Asian C. episcopus Woollyneck Storks. SIS Conserv. 80–95.

Gutiérrez, E., and K. Helgen. 2013. Outdated taxonomy blocks conservation. Nature 495:314.

Hall, R. 2009. Southeast Asia’s changing palaeogeography. Blumea - Biodivers. Evol. Biogeogr. Plants 54:148–161.

Harvey, M. G., G. A. Bravo, S. Claramunt, A. M. Cuervo, G. E. Derryberry, J. Battilana, G. F. Seeholzer, J. S. McKay, B. C. O’Meara, and B. C. Faircloth. 2020. The evolution of a tropical biodiversity hotspot. Science 370:1343–1348. American Association for the Advancement of Science.

Hasan, M., and P. Ghimire. 2020. Confirmed breeding records of Asian Woollyneck Ciconia episcopus from Bangladesh. SIS Conserv. 47–49.

Hernández-Romero, P. C., C. Gutiérrez-Rodríguez, C. Valdespino, and D. A. Prieto-Torres. 2018. The Role of Geographical and Ecological Factors on Population Divergence of the Neotropical otter Lontra longicaudis (Carnivora, Mustelidae). Evol. Biol. 45:37–55.

Keith, S. A., A. H. Baird, T. P. Hughes, J. S. Madin, and S. R. Connolly. 2013. Faunal breaks and species composition of Indo-Pacific corals: the role of plate tectonics, environment and habitat distribution. Proc. R. Soc. B Biol. Sci. 280:20130818. Royal Society.

Kittur, S., and K. S. G. Sundar. 2020. Density, flock size and habitat preferance of Woolly-necked Storks Ciconia episcopus in agricultural landscapes of south Asia. SIS Conserv. 71–79.

Lamichhaney, S., J. Berglund, M. S. Almen, K. Maqbool, M. Grabherr, A. Martinez-Barrio, M. Promerova, C.-J. Rubin, C. Wang, N. Zamani, B. R. Grant, P. R. Grant, M. T. Webster, and L. Andersson. 2015. Evolution of Darwin’s finches and their beaks revealed by genome sequencing. Nature 518:371– 375.

Li, H., and R. Durbin. 2009. Fast and accurate short read alignment with Burrows–Wheeler transform. Bioinformatics 25:1754–1760.

Liu, S., Y. Liu, E. Jelen, M. Alibadian, C.-T. Yao, X. Li, N. Kayvanfar, Y. Wang, F. S. M. Vahidi, J.-L. Han, G. Sundev, Z. Zhang, and M. Schweizer. 2020. Regional drivers of diversification in the late Quaternary in a widely distributed generalist species, the common pheasant Phasianus colchicus. J. Biogeogr. 47:2714–2727.

Long, J. A. 2017. Why Australasian vertebrate animals are so unique – A palaeontological perspective. Gen. Comp. Endocrinol. 244:2–10.

Malinsky, M., M. Matschiner, and H. Svardal. 2021. Dsuite - Fast D-statistics and related admixture evidence from VCF files. Mol. Ecol. Resour. 21:584–595. John Wiley & Sons, Ltd.

Mayr, E. 1982. Speciation and Macroevolution. Evolution 36:1119–1132. [Society for the Study of Evolution, Wiley].

Mayr, E. 1999. Systematics and the origin of species, from the viewpoint of a zoologist. Harvard University Press.

McIntyre, S. R. N., C. H. Lineweaver, C. P. Groves, and A. Chopra. 2017. Global biogeography since Pangaea. Proc. R. Soc. B Biol. Sci. 284:20170716. Royal Society.

McKenna, A., M. Hanna, E. Banks, A. Sivachenko, K. Cibulskis, A. Kernytsky, K. Garimella, D. Altshuler, S. Gabriel, M. Daly, and M. A. DePristo. 2010. The Genome Analysis Toolkit: A MapReduce framework for analyzing next-generation DNA sequencing data. Genome Res. 20:1297–1303.

McQuarrie, N., and D. J. J. van Hinsbergen. 2013. Retrodeforming the Arabia-Eurasia collision zone: Age of collision versus magnitude of continental subduction. Geology 41:315–318.

Mlodinow, S. G., J. del Hoyo, A. Elliott, N. Collar, E. F. J. Garcia, P. F. D. Boesman, and G. M. Kirwan. 2022. Asian Woolly-necked Stork (Ciconia episcopus), version 1.1. In Birds of the World (S. M. Billerman and B. K. Keeney, Editors). P. in Birds of the World. Cornell Lab of Ornithology, Ithaca, NY, USA.

National Academies of Sciences, Engineering, and Medicine and National Academies of Sciences, Engineering, and Medicine. 2019. Evaluating the Taxonomic Status of the Mexican Gray Wolf and the Red Wolf. The National Academies Press, Washington, DC.

Orlando, L., R. Allaby, P. Skoglund, C. Der Sarkissian, P. W. Stockhammer, M. C. Ávila-Arcos, Q. Fu, J. Krause, E. Willerslev, A. C. Stone, and C. Warinner. 2021. Ancient DNA analysis. Nat. Rev. Methods Primer 1:14.

Padial, J. M., A. Miralles, I. De la Riva, and M. Vences. 2010. The integrative future of taxonomy. Front. Zool. 7:16.

Pellissier, L., C. Heine, D. F. Rosauer, and C. Albouy. 2018. Are global hotspots of endemic richness shaped by plate tectonics? Biol. J. Linn. Soc. 123:247–261.

Pickrell, J. K., and J. K. Pritchard. 2012. Inference of Population Splits and Mixtures from Genome-Wide Allele Frequency Data. PLOS Genet. 8:e1002967. Public Library of Science.

Price, M. N., P. S. Dehal, and A. P. Arkin. 2010. FastTree 2 – approximately maximum-likelihood trees for large alignments. PLoS ONE 5:e9490–e9490. Public Library of Science.

Rannala, B. 2015. The art and science of species delimitation. Curr. Zool. 61:846–853.

Ravinet, M., R. Faria, R. K. Butlin, J. Galindo, N. Bierne, M. Rafajlović, M. A. F. Noor, B. Mehlig, and A. M. Westram. 2017. Interpreting the genomic landscape of speciation: a road map for finding barriers to gene flow. J. Evol. Biol. 30:1450–1477. John Wiley & Sons, Ltd.

Schluter, D. 2001. Ecology and the origin of species. Trends Ecol. Evol. 16:372–380. Elsevier.

Schluter, D. 2009. Evidence for ecological speciation and its alternative. Science 323:737–741. American Association for the Advancement of Science.

Smith, M. L., and B. C. Carstens. 2022. Species Delimitation Using Molecular Data. Pp. 145–160 in Species Problems and Beyond. CRC Press, Boca Raton.

Sobel, J. M., G. F. Chen, L. R. Watt, and D. W. Schemske. 2010. The biology of speciation. Evolution 64:295–315. Blackwell Publishing Inc Malden, USA.

Sundar, K. G. 2020. Woolly-necked Stork -a species ignored. 2:33–41.

Team, R. C. 2016. R: A language and environment for statistical computing. R Foundation for Statistical Computing, Vienna, Austria. Http://www.R-Proj.Org.

Thabethe, V., and C. T. Downs. 2018. Citizen science reveals widespread supplementary feeding of African woolly-necked storks in suburban areas of KwaZulu-Natal, South Africa. Urban Ecosyst. 21:965–973.

Tobias, J. A., N. Seddon, C. N. Spottiswoode, J. D. Pilgrim, L. D. C. Fishpool, and N. J. Collar. 2010. Quantitative criteria for species delimitation. Ibis 152:724–746.

Tremble, K., J. I. Hoffman, and B. T. M. Dentinger. 2023. Contrasting continental patterns of adaptive population divergence in the holarctic ectomycorrhizal fungus Boletus edulis. New Phytol. 237:295–309.

Vaghela, U., D. Sawant, and V. Bhagwat. 2015. Woolly-necked Storks Ciconia episcopus nesting on mobile-towers in Pune, Maharashtra. Indian BIRDS 10:154–155.

Van der Auwera, G. A., M. O. Carneiro, C. Hartl, R. Poplin, G. Del Angel, A. Levy-Moonshine, T. Jordan, K. Shakir, D. Roazen, J. Thibault, E. Banks, K. V. Garimella, D. Altshuler, S. Gabriel, and M. A. DePristo. 2013. From FastQ data to high confidence variant calls: the Genome Analysis Toolkit best practices pipeline. Curr. Protoc. Bioinforma. 43:11.10.1–33.

Verstappen, H. Th. 1997. The effect of climatic change on southeast Asian geomorphology. J. Quat. Sci. 12:413–418.

Yuan, Z.-Y., B.-L. Zhang, C. J. Raxworthy, D. W. Weisrock, P. M. Hime, J.-Q. Jin, E. M. Lemmon, A. R. Lemmon, S. D. Holland, M. L. Kortyna, W.-W. Zhou, M.-S. Peng, J. Che, and E. Prendini. 2019. Natatanuran frogs used the Indian Plate to step-stone disperse and radiate across the Indian Ocean. Natl. Sci. Rev. 6:10–14.

Zachos, F. E. 2013. Species splitting puts conservation at risk. Nature 494:35–35. Nature Publishing Group.

Zapata, F., and I. Jiménez. 2012. Species delimitation: inferring gaps in morphology across geography. Syst. Biol. 61:179. Oxford University Press.

