## Supplementary Fig. 1, Supplementary Fig. 2, Supplementary Fig. 3, Supplementary Fig. 4, Supplementary Fig. 5, Supplementary Table 1 for "Unraveling cryptic diversity: Genomic approaches to study the taxonomy and evolution of Woolly-necked storks using museum specimens"

**Supplementary Figures**

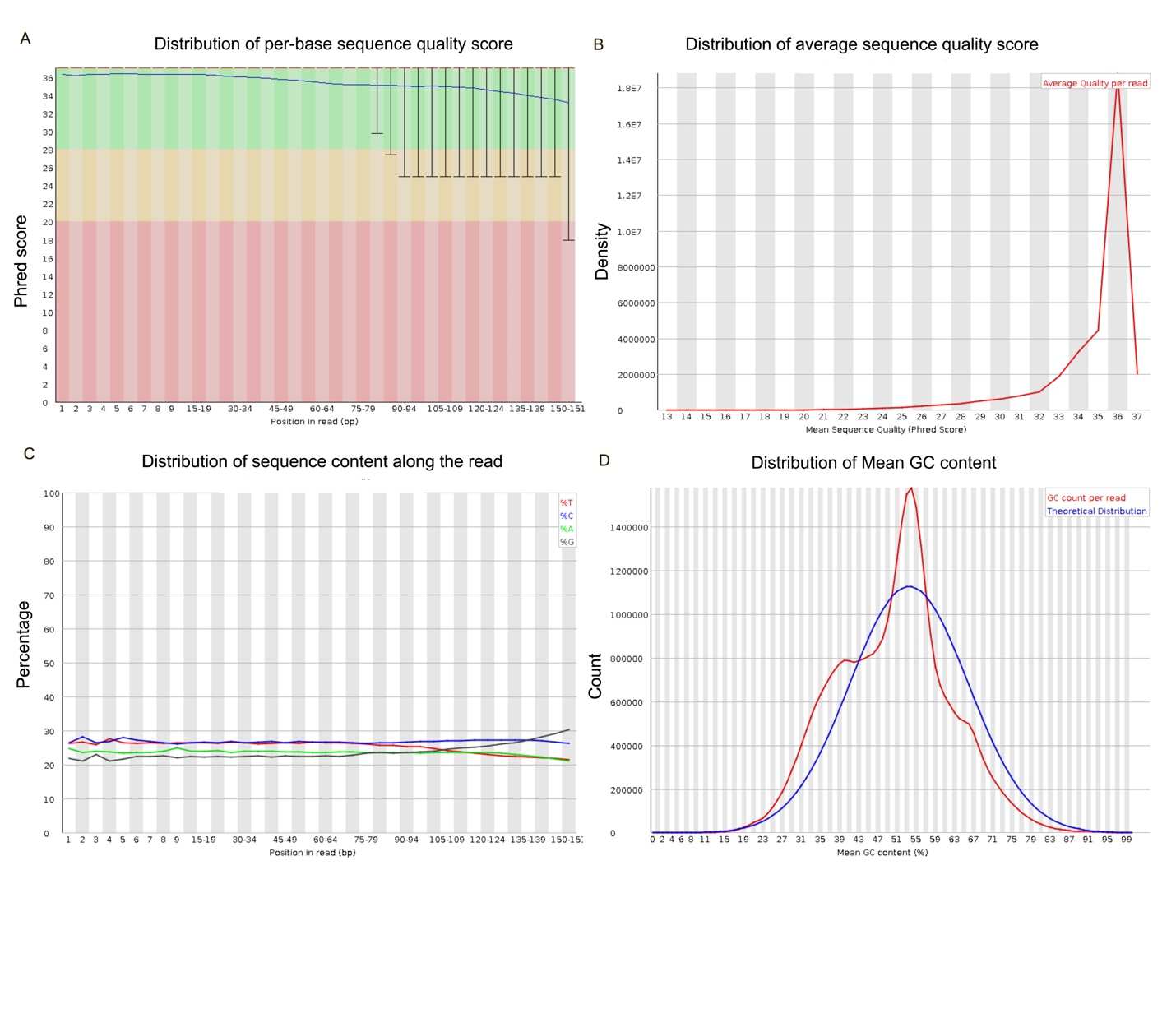
**Supplementary Fig. 1:** FASTQC report from sample MCZ:264670 (*Ciconia microscelis*) (A) Distribution of per-base sequence quality score (B) Distribution of average sequence quality score (C) Distribution of sequence content along the read (D) Distribution of Mean GC content. The quality report was similar for all 12 samples.

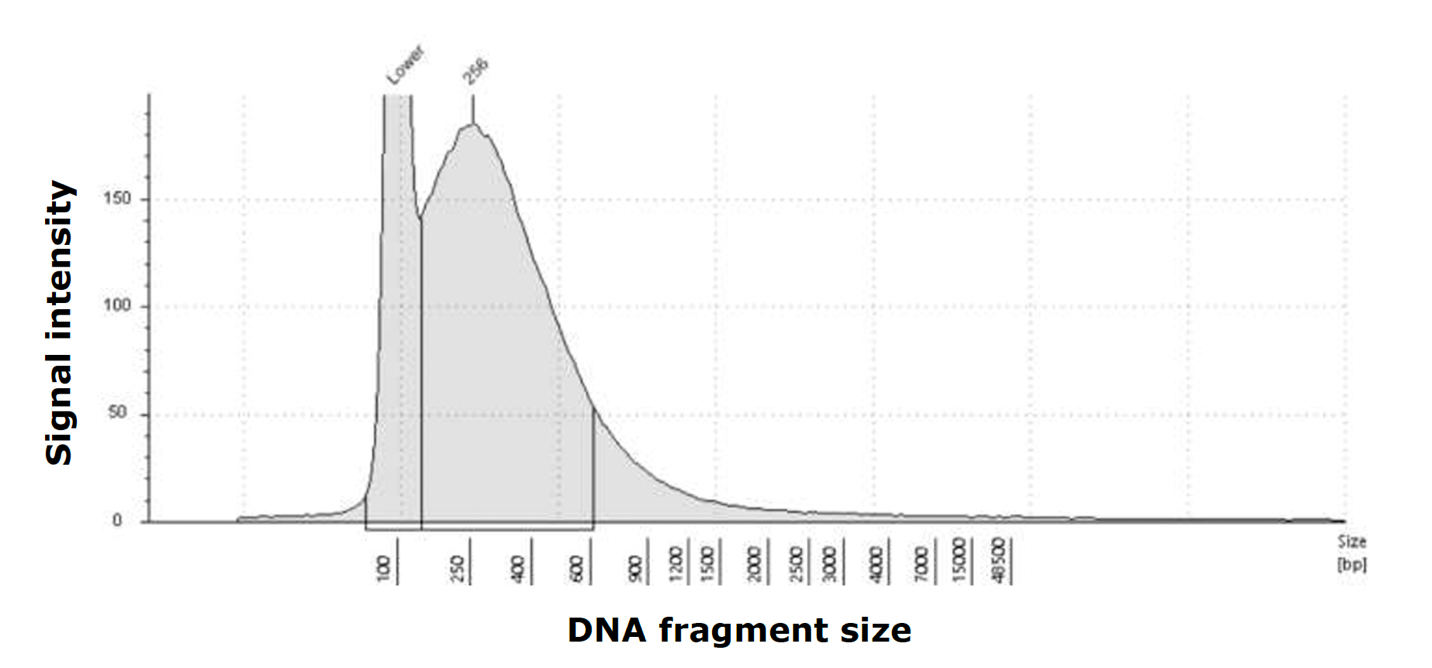

**Supplementary Fig. 2:** Distribution of fragment lengths of genomic DNA extracted from sample MCZ:264670 (Ciconia microscelis). Great majority of DNA had shorter fragments indicative of DNA degradation. The distribution of fragment lengths was similar for all 12 samples.

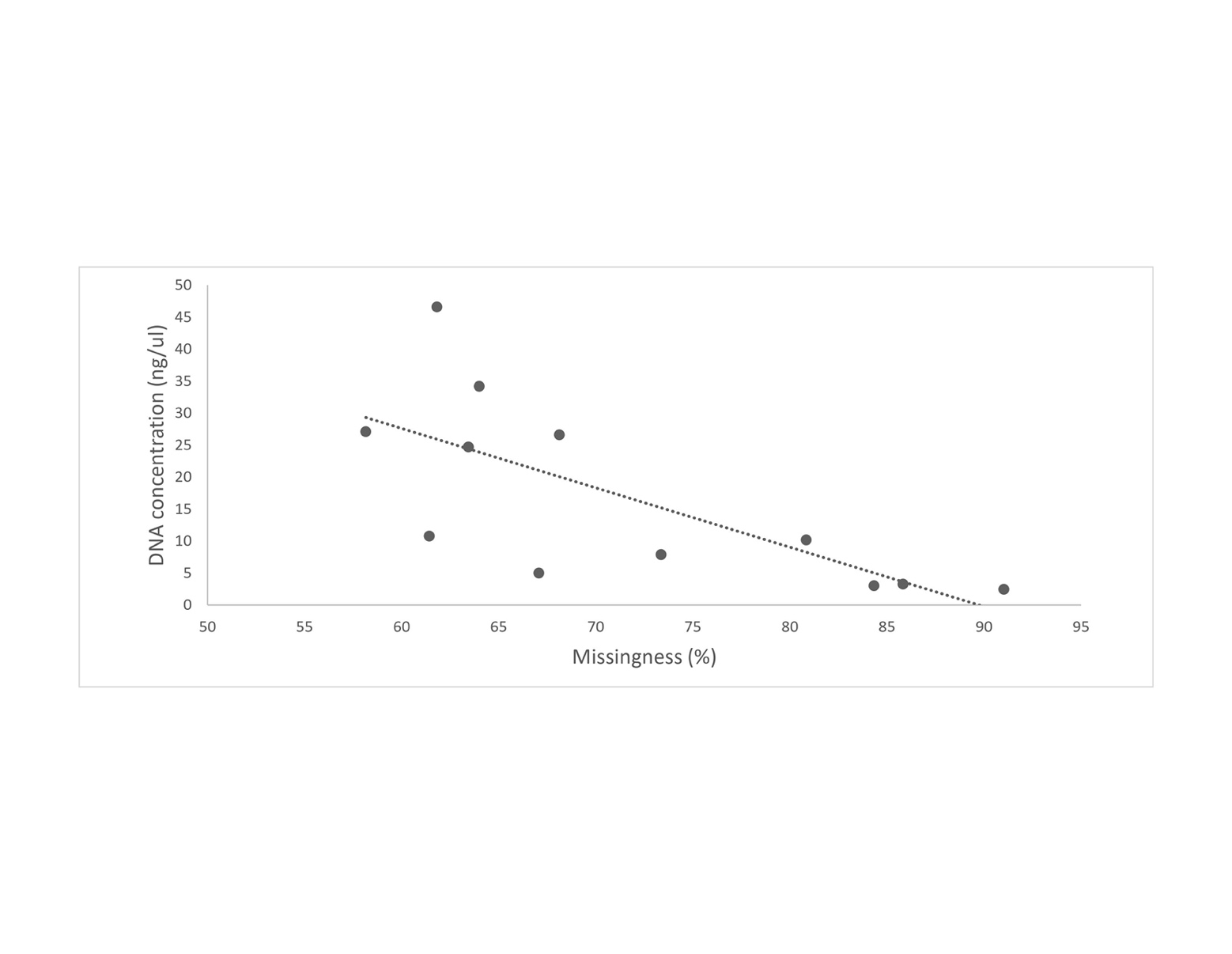

**Supplementary Fig. 3**: Correlation between DNA quality (measured as DNA concentration) and percentage of missing SNPs across all samples. Lower quality DNA had higher rate of missing SNPs.

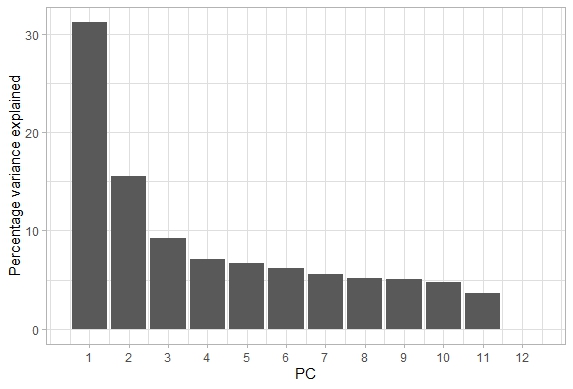

**Supplementary Fig. 4:** Percentage of variable explained by each 12 principal components (PCs). First two PCs explained almost 50% of genetic variations among samples.

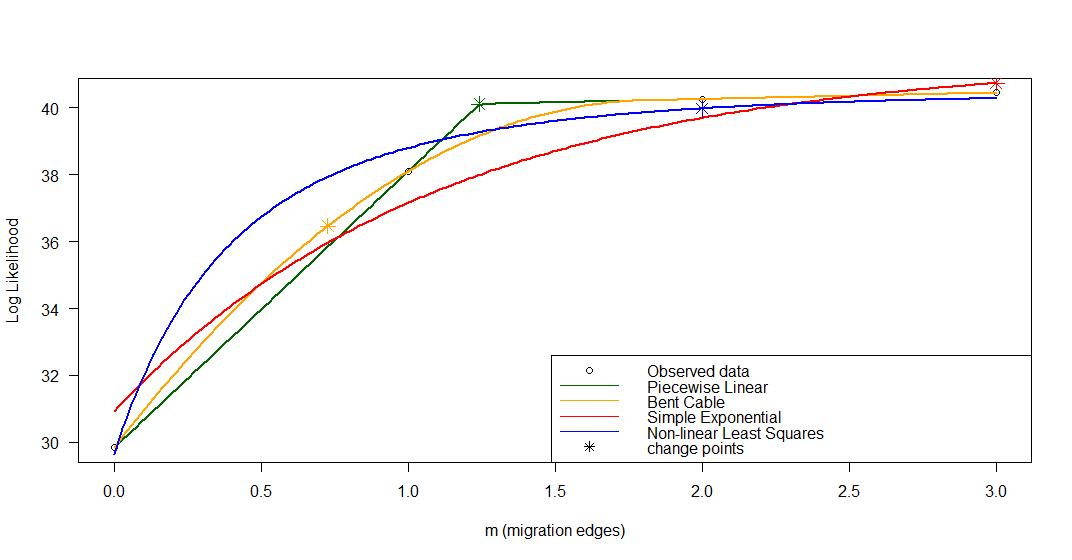

**Supplementary Fig. 5:** Plot showing changes in log-likelihood for a possible number of migration events (m). Values produced after 500 bootstraps. Migration event of either 1 or 2 appears optimum.

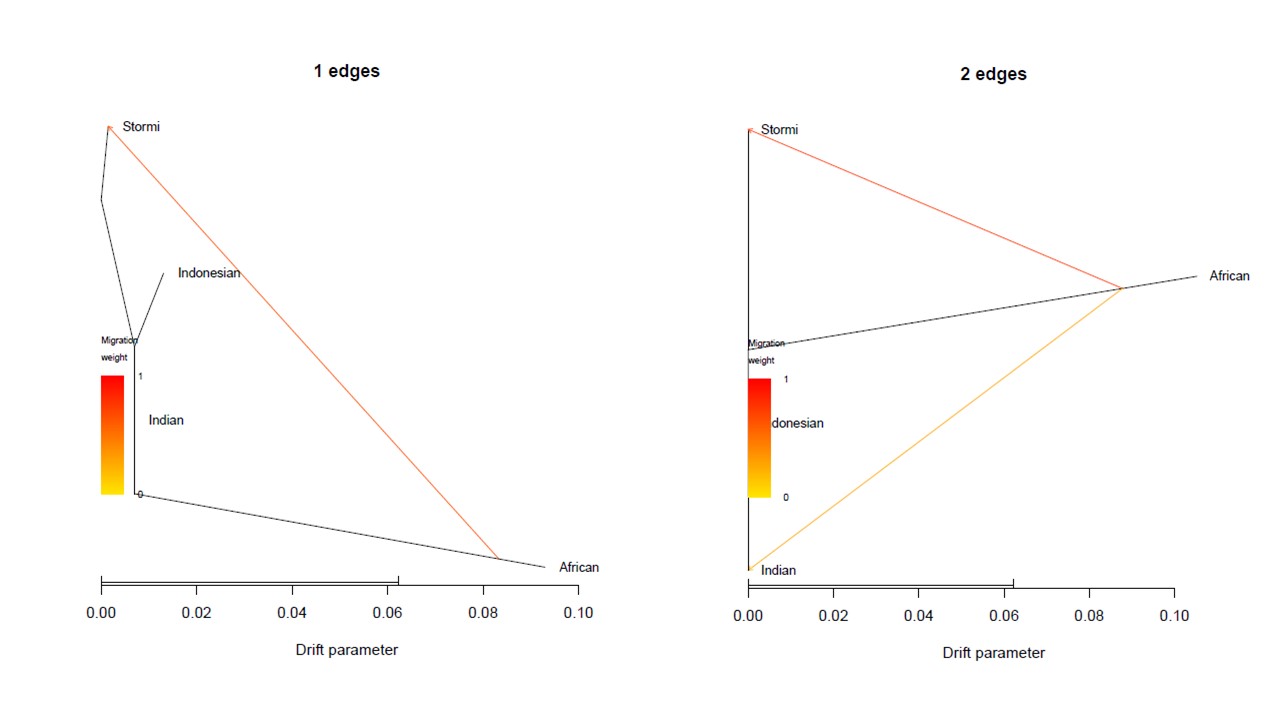

**Supplementary Fig. 6:** Population graphs inferred by TreeMix (Pickrell & Pritchard, 2012) using 33,087 genome-wide SNPs for m=1 and m = 2 edges. Branch lengths are proportional to the evolutionary change (the drift parameter) and terminal nodes are labeled with population codes (see Supplementary Table 1 for details).

**Supplementary Table 1:** Details of museum specimen used in this study.

| **Species** | **MCZ^*^ Catalogue Number** | **Sampling location** | **Sampling Date** |
| --- | --- | --- | --- |
| *Ciconia microscelis* | 11616 | South Africa | NA^**^ |
| *Ciconia microscelis* | 12831 | African Origin | NA^**^ |
| *Ciconia microscelis* | 264670 | Liberia | 1939 |
| *Ciconia microscelis* | 270607 | Ganta via Monrovia, Tanzania | 1939 |
| *Ciconia microscelis* | 279738 | Miranja , Sierra Leone | 1953 |
| *Ciconia stormi* | 170531 | Nongoba Bullom Chiefdom, Indonesia | 1934 |
| *Ciconia stormi* | 170532 | West Sumatra, Indonesia | 1934 |
| *Ciconia episcopus neglecta* | 270046 | West Sumatra, Indonesia | 1939 |
| *Ciconia episcopus neglecta* | 270047 | Sulawesi Island, Indonesia | 1939 |
| *Ciconia episcopus neglecta* | 270321 | Sulawesi Island, Indonesia | 1938 |
| *Ciconia episcopus episcopus* | 278124 | India | 1948 |
| *Ciconia episcopus episcopus* | 54164 | Unknown | 1953 |

*Museum of Comparative Zoology, Harvard University

**Not available

**Supplementary Table 2:** Total amount of DNA extracted from toe-pad skins samples of historic museum specimens used in this study.

| **Species** | **MCZ^*^ Catalogue Number** | **Total amount of DNA extracted (ng)** | **Total number of sequencing reads** | **Total genomic data (GB)** |
| --- | --- | --- | --- | --- |
| *Ciconia microscelis* | 11616 | 276 | 88,042,788 | 13.21 |
| *Ciconia microscelis* | 12831 | 294 | 65,064,226 | 9.76 |
| *Ciconia microscelis* | 264670 | 321.9 | 89,185,542 | 13.38 |
| *Ciconia microscelis* | 270607 | 429 | 77,001,948 | 11.55 |
| *Ciconia microscelis* | 279738 | 156 | 87,657,360 | 13.15 |
| *Ciconia stormi* | 170531 | 210 | 69,981,572 | 10.50 |
| *Ciconia stormi* | 170532 | 124 | 73,581,904 | 11.04 |
| *Ciconia episcopus neglecta* | 270046 | 126.3 | 77,323,416 | 11.60 |
| *Ciconia episcopus neglecta* | 270047 | 162 | 86,747,950 | 13.01 |
| *Ciconia episcopus neglecta* | 270321 | 210 | 81,744,096 | 12.26 |
| *Ciconia episcopus episcopus* | 278124 | 225 | 78,198,934 | 11.73 |
| *Ciconia episcopus episcopus* | 54164 | 327 | 84,796,218 | 12.72 |

**Supplementary Table 3**: Sequence alignment statistics by mapping short reads against the reference genome of Maguari stork (Ciconia maguari)

| **Species** | **MCZ^*^ Catalogue Number** | **Total sequence mapped to reference genome (%)** | **Both pair mapped to reference genome (%)** | **Average genome wide sequencing depth** |
| --- | --- | --- | --- | --- |
| *Ciconia microscelis* | 11616 | 92.33 | 84.46 | 5.69 |
| *Ciconia microscelis* | 12831 | 90.51 | 81.97 | 5.24 |
| *Ciconia microscelis* | 264670 | 98.91 | 89.96 | 6.63 |
| *Ciconia microscelis* | 270607 | 98.76 | 88.63 | 6.21 |
| *Ciconia microscelis* | 279738 | 99.12 | 85.55 | 6.16 |
| *Ciconia stormi* | 170531 | 83.2 | 76.68 | 4.5 |
| *Ciconia stormi* | 170532 | 97.54 | 91.3 | 5.59 |
| *Ciconia episcopus neglecta* | 270046 | 98.8 | 86.52 | 5.51 |
| *Ciconia episcopus neglecta* | 270047 | 98.62 | 90.25 | 6.86 |
| *Ciconia episcopus neglecta* | 270321 | 96.49 | 87.22 | 6.46 |
| *Ciconia episcopus episcopus* | 278124 | 99.34 | 87.59 | 6.23 |
| *Ciconia episcopus episcopus* | 54164 | 93.25 | 84.77 | 5.92 |

**Supplementary Table 4:** Statistics of missing SNPs (Missingness score = 0, SNP genotyped across all 12 samples; and score = 1, SNP missing in all samples)

| **Missingness Score** | **Total number of SNPs** | **% of total SNPs** |
| --- | --- | --- |
| < 0.1 | 131,965 | 0.97 |
| 0.1-0.2 | 224,543 | 1.66 |
| 0.2-0.3 | 439,916 | 3.25 |
| 0.3-0.4 | 770,401 | 5.69 |
| 0.4-0.5 | 1,216,984 | 8.98 |
| 0.5-0.6 | 4,129,831 | 30.48 |
| >0.7 | 6,636,116 | 48.98 |

**Supplementary Table 5:** Cross-validation error (CV) for k = 1–5 ancestry (K) shows that k= 3 or 4 are the optimal number of genetic clusters for admixture analysis.

| **Ancestry(k)** | **Cross-validation error (CVV)** |
| --- | --- |
| 1 | 0.51043 |
| 2 | 0.65317 |
| 3 | 0.53515 |
| 4 | 0.53604 |
| 5 | 0.60235 |

**Supplementary Table 6:** Estimates of ABBA-BABA analysis using Dsuite (Malinsky et al. 2021) to identify evidence of gene flow between two clades of Asian Wollynecks (neglecta and episcopus) (P1 and P2) and African Wollynecks (P3) using Storm’s stork as an outgroup.

| **P1** | **P2** | **P3** | **Dstatistics** | **Z-score** | **p-value** | **f4-ratio** | **BBAA** | **ABBA** | **BABA** |
| --- | --- | --- | --- | --- | --- | --- | --- | --- | --- |
| *Ciconia episcopus neglecta* | *Ciconia episcopus episcopus* | *Ciconia microscelis* | 0.014 | 1.42 | 0.07 | 0.029 | 1156.02 | 648.246 | 629.854 |
| *Ciconia episcopus episcopus* | *Ciconia microscelis (clade 1: Tanzania)* | *Ciconia microscelis (clade 2)* | 0.324 | 9.246 | 0.00 | 0.862 | 599.208 | 1206.4 | 615.312 |
